# Mutually opposing activity of PIN7 splicing isoforms is required for auxin-mediated tropic responses in *Arabidopsis thaliana*

**DOI:** 10.1101/2020.05.02.074070

**Authors:** Ivan Kashkan, Mónika Hrtyan, Katarzyna Retzer, Jana Humpolíčková, Aswathy Jayasree, Roberta Filepová, Zuzana Vondráková, Sibu Simon, Debbie Rombaut, Thomas B. Jacobs, Mikko J. Frilander, Jan Hejátko, Jiří Friml, Jan Petrášek, Kamil Růžička

**Affiliations:** Laboratory of Hormonal Regulations in Plants, Institute of Experimental Botany, Czech Academy of Sciences, 16502 Prague, Czech Republic; Functional Genomics and Proteomics of Plants, Central European Institute of Technology and National Centre for Biomolecular Research, Masaryk University, 62500 Brno, Czech Republic; Institute of Organic Chemistry and Biochemistry, Czech Academy of Sciences, 166 10 Prague 6, Czech Republic; Department of Plant Biotechnology and Bioinformatics, Ghent University, 9052 Ghent, Belgium; VIB Center for Plant Systems Biology, 9052 Ghent, Belgium; Institute of Biotechnology, University of Helsinki, 00014 University of Helsinki, Finland; Institute of Science and Technology (IST Austria), 3400 Klosterneuburg, Austria

**Keywords:** Auxin, auxin transport, PINs, alternative splicing, plant development, RNA processing, tropic responses

## Abstract

Advanced transcriptome sequencing has uncovered that the majority of eukaryotic genes undergo alternative splicing (AS). Nonetheless, little effort has been dedicated to investigating the functional relevance of particular splicing events, even those in the key developmental and hormonal regulators. Here we reveal, in the plant model *Arabidopsis thaliana*, that the *PIN7* gene, which encodes a polarly localized transporter for the phytohormone auxin, produces two evolutionary-conserved transcripts. These isoforms PIN7a and PIN7b, differing in a 4 amino acid motif, are present at nearly equal levels in most cells. Although both variants do not differ in the subcellular localization and transport auxin with similar capacity, they closely associate and mutually influence their stability within the plasma membrane. Phenotypic complementation tests reveal that the functional contribution of PIN7b *per se* is minor but it markedly attenuates the prominent PIN7a activity, which is required for correct seedling apical hook formation and auxin-mediated tropic responses. These results establish alternative splicing of the PIN family as a conserved, functionally-relevant mechanism, unveiling an additional regulatory level of auxin-mediated plant development.

## INTRODUCTION

Auxin is an essential phytohormone, which plays a role in nearly all aspects of plant development. To flexibly adapt to rapidly-changing environmental cues, directional auxin transport represents a highly dynamic means for triggering downstream morphogenetic processes. PIN-FORMED (PIN) auxin efflux carriers are among the key regulators in this respect. In the past years, many efforts uncovered multiple mechanisms operating transcriptionally or post-translationally on the capacity and directionality of PIN-mediated transport. However, little progress has been made in exploring the contribution of post-transcriptional regulation (Hrtyan et al., 2015; Adamowski and Friml, 2015).

Advances in high throughput sequencing have revealed unexpected complexity within eukaryotic transcriptomes by alternative splicing (AS). Although the majority of AS transcripts may be functionally neutral (Tress et al., 2017; Blencowe, 2017; Darracq and Adams, 2013; Chamala et al., 2015; Reddy et al., 2013), several detailed studies have highlighted a plausible role for numerous AS events in physiologically relevant contexts, including those involved in plant developmental and hormonal pathways (Shang et al., 2017; Hrtyan et al., 2015; Staiger and Brown, 2013; Szakonyi and Duque, 2018). Earlier works have described auxin-related defects resulting from the aberrant function of several regulators of AS (Hrtyan et al., 2015; Kalyna et al., 2003; Tsugeki et al., 2015; Casson et al., 2009; Retzer et al., 2014; Bazin et al., 2018). AS changes subcellular localization of the auxin biosynthetic gene *YUCCA 4* (Kriechbaumer et al., 2012) and differential splicing of an exitron (Marquez et al., 2015) inside the *AUXIN RESPONSE FACTOR 8* (*ARF8*) results in developmental changes of generative organs (Ghelli et al., 2018). Likewise, a splice variant of *MONOPTEROS* (*MP*/*ARF5*) acts as a noncanonical regulator of ovule development (Cucinotta et al., 2020). In addition, AS of the Major Facilitator Superfamily transporter ZIFL1 interferes with auxin transport, influencing the stability of PINs on the plasma membrane (PM) (Remy et al., 2013). These lines of evidence indicate that AS is an important player in auxin-dependent processes. However, no coherent functional model of any auxin-related AS event has been provided so far.

Here, we have investigated the functional significance of the protein isoforms arising through AS of the *Arabidopsis thaliana PIN7* gene. PIN7 is, together with PIN3 and PIN4, a member of the PIN3 clade of PIN auxin efflux carriers (Bennett et al., 2014), which are required for a broad range of morphogenetic and tropic processes (Adamowski and Friml, 2015). We provide evidence that the evolutionary conserved PIN7 isoforms mutually influence their dynamics within PM and demonstrate that the coordinated action of both splice variants is required for auxin-mediated differential growth responses.

## RESULTS

### *Arabidopsis PIN7* and *PIN4* produce two evolutionary conserved AS transcripts

Our previous survey (Hrtyan et al., 2015) revealed that several genes involved in auxin-dependent processes undergo AS. Among them, closely related paralogs from the *PIN3* clade of auxin transporters, *PIN4* and *PIN7* (but not *PIN3*), are regulated by the same type of AS (Petrasek et al., 2006; Bennett et al., 2014; Hrtyan et al., 2015). The resulting transcripts, denoted as *a* and *b*, differ in the AS donor site position in the first intron (Figure 1A). The differentially spliced region corresponds to a four amino acid motif inside the large internal hydrophilic loop (Nodzyński et al., 2016; Ganguly et al., 2014) of the integral PM transporter (Figure 1A and 1B). We examined the quantities of individual reads spanning the exon junctions in the respective gene region from the *Arabidopsis* root tip and in several other available transcriptomes from different tissues and organs (Cheng et al., 2017; Klepikova et al., 2016; Ruzicka et al., 2017) (Figures 1C and Supplemental Table 1). We found that each of the *PIN4* and *PIN7* splice isoforms is expressed abundantly in all tissues, independently of the inspected data set. We also identified a minor *PIN4c* (Hrtyan et al., 2015; Marquez et al., 2012) transcript (but not corresponding *PIN7c*), which comprised around 3-7% of the *PIN4* exon1-exon2 spanning reads (Figures 1A and Supplemental Table 1). Other occasionally observed transcripts (also corresponding to the other exon junctions) were not seen among different RNA-seq data sets. It thereby appears that *PIN7* and *PIN4* are processed into two and three splice isoforms, respectively, and that *PIN7a* and *b* (or *PIN4a* and *b*) are expressed in most of the plant organs at comparable or nearly similar levels.

**Figure 1.**
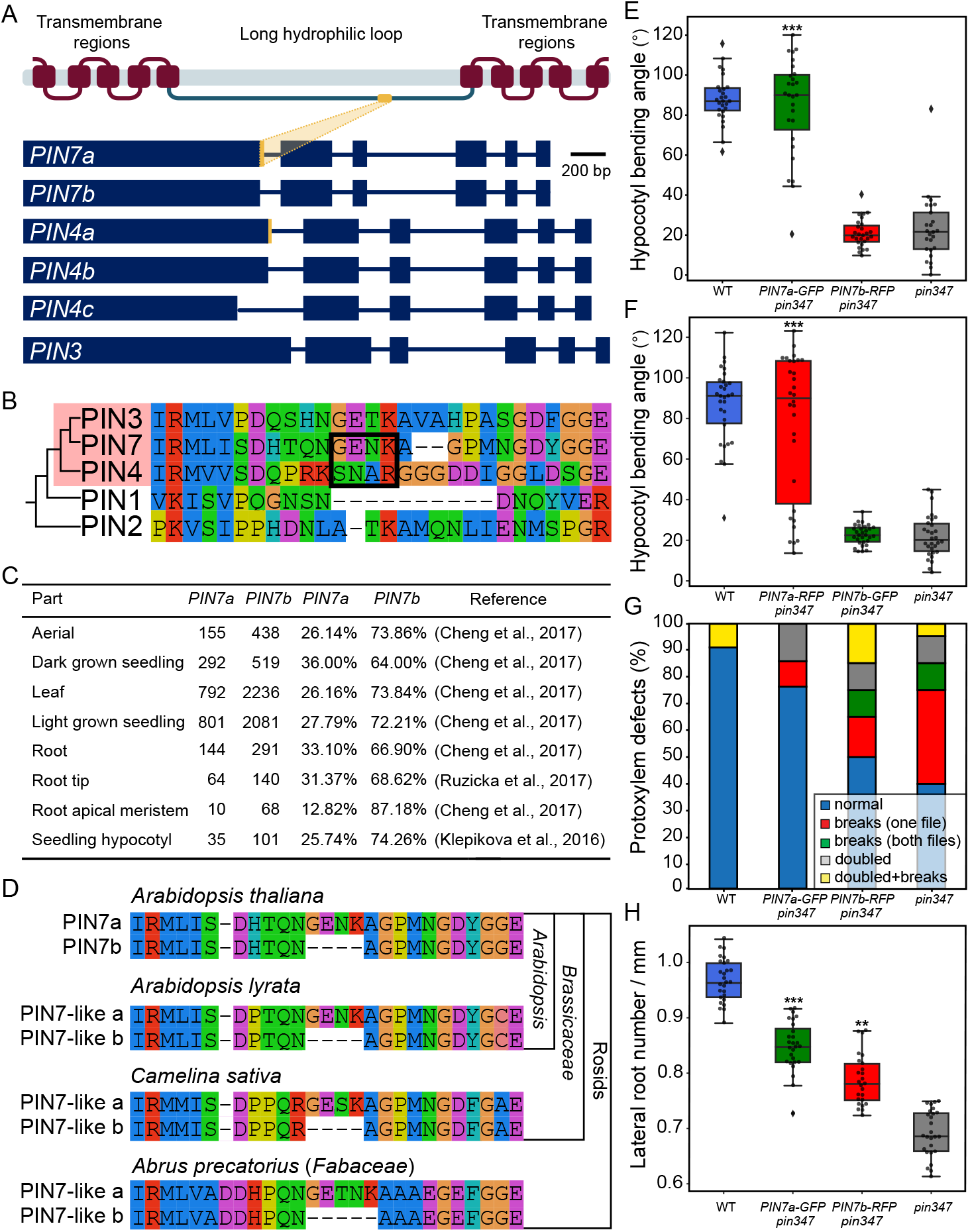
Alternative splicing (AS) of the *Arabidopsis thaliana PIN7* and *PIN4* genes leads to two evolutionary conserved functionally different transcripts. **(A)**Scheme of coding regions of the *PIN3* clade genes in *Arabidopsis thaliana*. The alternative donor splice site at the end of the first exon of *PIN7* and *PIN4*, respectively, but not *PIN3*, results in two transcripts differing in 12 nucleotides. This sequence (orange) corresponds to the protein motif, located in the long internal hydrophilic loop of the transporter. There is also an additional *PIN4c* transcript present in publicly available transcriptomes. **(B)**Amino acid alignment of the region around the 4-amino acid motif changed by AS in the PIN3 clade proteins (boxed in pink) in *Arabidopsis thaliana*, including their closest PIN paralogs. **(C)**Table shows the number of RNA-seq reads spanning the exon1-exon2 junction corresponding to the detected *PIN7* transcripts in selected *Arabidopsis thaliana* tissue sources. Their ratio was calculated as a percentage of total reads mapped to this area, as assessed from the genome browser graphic interface. **(D)** Protein sequence alignments showing the conservation of AS in the PIN3 clade of auxin transporters among rosids. **(E)**and **(F)** Phototropic bending of the etiolated *pin347* seedlings carrying the *PIN7a-GFP* and *PIN7b-RFP* **(E)**, and the *PIN7a-RFP* and *PIN7b-GFP* constructs **(F). (G)** and **(H)** Quantification of the primary root protoxylem defects **(G)** and lateral root primordia **(H)** initiation of the *pin347* seedlings harboring the *PIN7a-RFP* and *PIN7b-GFP* constructs. On **(E), (F)** and **(H)**, the middle line corresponds to median, the box corresponds to the 25% and 75% quantiles, the whiskers are the minima and maxima, dots represent single data points. The asterisks indicate a difference between the respective line and the *pin347* mutant (\*\**P* < 0.01, \*\*\**P* < 0.001 by two-way ANOVA). For each line in each experiment, at least 15 seedlings were analyzed.

Functionally relevant AS events are commonly evolutionary conserved (Reddy et al., 2013; Keren et al., 2010), therefore we sought for available validated transcripts to determine whether similar splicing events occur in orthologous *PIN3* clade genes in other dicot species (Bennett et al., 2014; O ‘Leary et al., 2016). We found examples of such mRNAs besides members of the *Brassicaceae* family, for instance, in *Abrus precatorius* (*Fabaceae*), which documents the conservation of this AS event in rosids, a plant clade that diversified more than 100 million years ago (Li et al., 2019) (Figure 1D). *PIN4c* did not show any deeper evolutional conservation. Thus, at least some genes of the *PIN3* clade are regulated by the same type of AS, across several plant families, which suggests that these AS events may have a relevant biological function.

### *PIN7* splice isoforms are functionally distinct, based on the mutant phenotype rescue tests

We expressed fluorescently-tagged cDNA versions of the respective *PIN7* and *PIN4* transcripts under their native promoters in *Arabidopsis thaliana*. Their overall expression patterns resembled those of the *PIN7-GFP* and *PIN4-GFP* lines made on the basis of the genomic sequence (Supplemental Figures 1A –1D). Next, we tested their ability to complement the phenotypes associated with the *PIN7* locus. The phototropic hypocotyl bending is among the classical assays for testing the activity of the PIN3 clade proteins (Friml et al., 2002; Willige et al., 2013). As weak phenotypes of the *pin7-2* loss of function mutants are the result of redundancy with other genes from the *PIN3* clade (Blilou et al., 2005; Willige et al., 2013; Friml et al., 2003), we employed a triple *pin3-3 pin4-101 pin7-102* knock out (*pin347*) as a genetic background, which lacks the phototropic response almost completely (Willige et al., 2013). Here, *PIN7a-GFP* cDNA rescued the phototropic bending, while the *PIN7b-RFP* cDNA did not show any effect, regardless of whether the native (Figure 1E) or a stronger endodermal *SCR* promoter (Rakusová et al., 2011) (Supplemental Figure 1E) was used. The tag choice does not appear to have any effect in these assays, as evidenced by the lines where the fluorophore sequences have been swapped (Figure 1F). Together, these data indicate that the motif changed by AS alters the function of the PIN7 protein in *Arabidopsis*.

The PIN3 clade auxin efflux carriers have been implicated in many other instances of auxin-mediated development (Adamowski and Friml, 2015). We therefore hypothesized whether, in some of them, the role of the particular isoform could be prevalent. These processes include: determining of root protoxylem formation (Bishopp et al., 2011) (Figure 1G), lateral root density (Swarup et al., 2008) (Figure 1H), vertical direction of the root growth (Kleine-Vehn et al., 2010; Friml et al., 2002; Pernisova et al., 2016) (Supplemental Figure 1F), lateral root orthogravitropism (Rosquete et al., 2013) (Supplemental Figure 1G), gravity-induced hypocotyl bending (Rakusová et al., 2011) (Supplemental Figure 1H), number of rosette branches after decapitation (Bennett et al., 2016) (Supplemental Figure 1I) and the overall rosette size (Bennett et al., 2016) (Supplemental Figure 1J). In sum, the *PIN7a-GFP* cDNA usually almost completely rescues the *pin347* phenotypes, while the contribution of *PIN7b-RFP* is smaller or even undetectable.

### Fluorescent reporters reveal highly overlapping *PIN7a* and *b* expression

The levels of *PIN7* (and *PIN4*) AS transcripts seem to be at comparable levels in most organs (Figure 1C and Supplemental Table 1). However, this may not describe the actual situation at the resolution of individual cells. To address this, we analyzed the P7A_1_G and P7BR fluorescent reporters (Kashkan et al., 2020), which allow for monitoring the activity of the AS of *PIN7 in planta* and *in situ* (Figure 2A). Indeed, in most cells, including the primary root tip (Figure 2B) or in the hypocotyl during the light bending assay (Figure 2C), we observed generally overlapping expression of both isoforms without any apparent tissue preference. However, there were several instances in the vegetative tissue, where the ratio of reporter signals appears to be uneven. These include early lateral root primordia (Supplemental Figure 1K), the mature pericycle adjacent to the phloem area (Supplemental Figure 1L), the stomatal lineage ground cells of the cotyledons (Supplemental Figure 1M), and the concave side of the opening apical hook (Figure 2D), where the PIN7b-RFP signal prevailed over that of PIN7a-GFP. In general, these data corroborate the presence of both isoforms in most cells and suggest that they may function in a coordinated manner.

**Figure 2.**
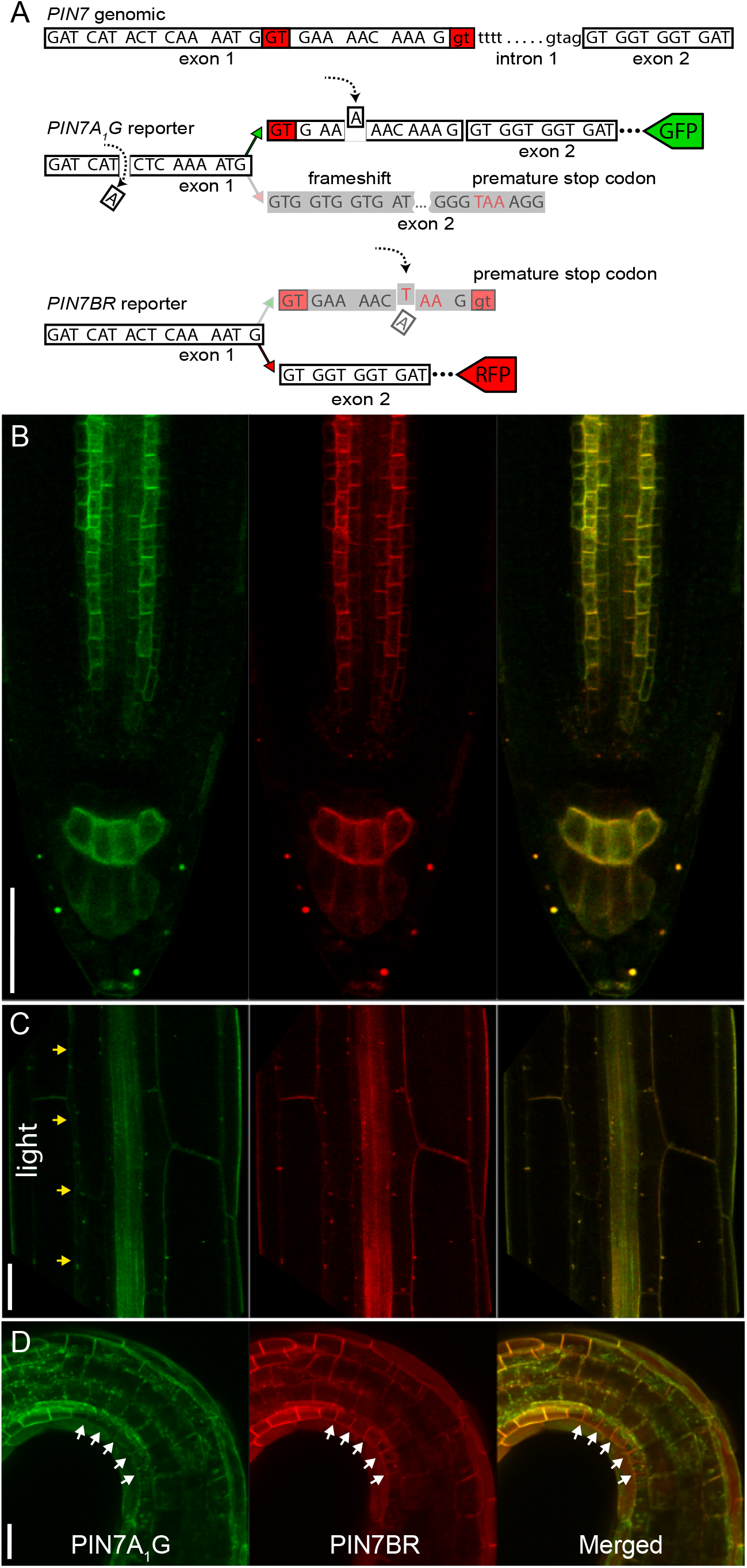
The PIN7 AS reporter system shows overlapping PIN7a and PIN7b expression in most tissues. **(A)** A scheme of the *PIN7* splicing reporter, which consists of two separate constructs. The *PIN7a-GFP* part (PIN7A_1_G) contains two point mutations (marked with dashed arrows) and leads to the frameshift and its subsequent restoration when the *PIN7a* transcript is solely produced. *PIN7b-RFP* reporter (PIN7BR) carries a premature stop codon inside the protruding *PIN7a* region (dashed arrow below). **(B)** to **(D)** Predominantly overlapping PIN7a-GFP and PIN7b-RFP reporter expression in the root tip **(B)**, in the etiolated hypocotyl following 4 h of unilateral light stimulation **(C)**, and in the area covering apical hook **(D)**. Yellow arrows: direction of unilateral light, white arrows: differential expression of the reporter. For each tissue, at least 9 images were analyzed. Bars, 50 µm.

### The combined activity of both PIN7a and PIN7b is required for apical hook formation and tropic responses

The generally overlapping expression and also an occasionally exaggerated response of the *pin347* mutants containing the *PIN7a-GFP* transgene (Supplemental Figure 1H) prompted us to carefully record the temporal dynamics of the processes linked with the PIN3 clade function. We were even able to capture subtle phenotypic changes observed in the single *pin7-2* knock out allele (Friml et al., 2003), including those complemented with the *PIN7* cDNAs during hypocotyl phototropism (Supplemental Figure 2A) or apical hook formation (Supplemental Figure 2B), where the time scale measurements of the PIN-mediated morphogenesis have been well established (Zadnikova et al., 2010). Next, we analyzed apical hook development in the *pin4-101 pin7-102* (*pin34*) mutants and a newly generated *pin347* line that carried a combination of both *PIN7* cDNA constructs. Similar to the gravi- and phototropic experiments, the *PIN7b-RFP* cDNA generally did not complement the severe *pin347* phenotype. Expression of the *PIN7a-GFP* cDNA in *pin347* indeed led to partial rescue of the phenotypic defects, even surpassing the values observed for *pin34*. Surprisingly, the simultaneous expression of both *PIN7a-GFP* and *PIN7b-RFP* suppressed the dominant effects conferred by the *PIN7a-GFP* cDNA alone and phenocopied the *pin34* mutant (Figure 3A; this effect was not caused by suppressing the expression of PIN7a-GFP by the other transgene, Supplemental Figure 2C). This strongly suggests that both isoforms act in a mutually opposing manner to modulate apical hook development.

**Figure 3.**
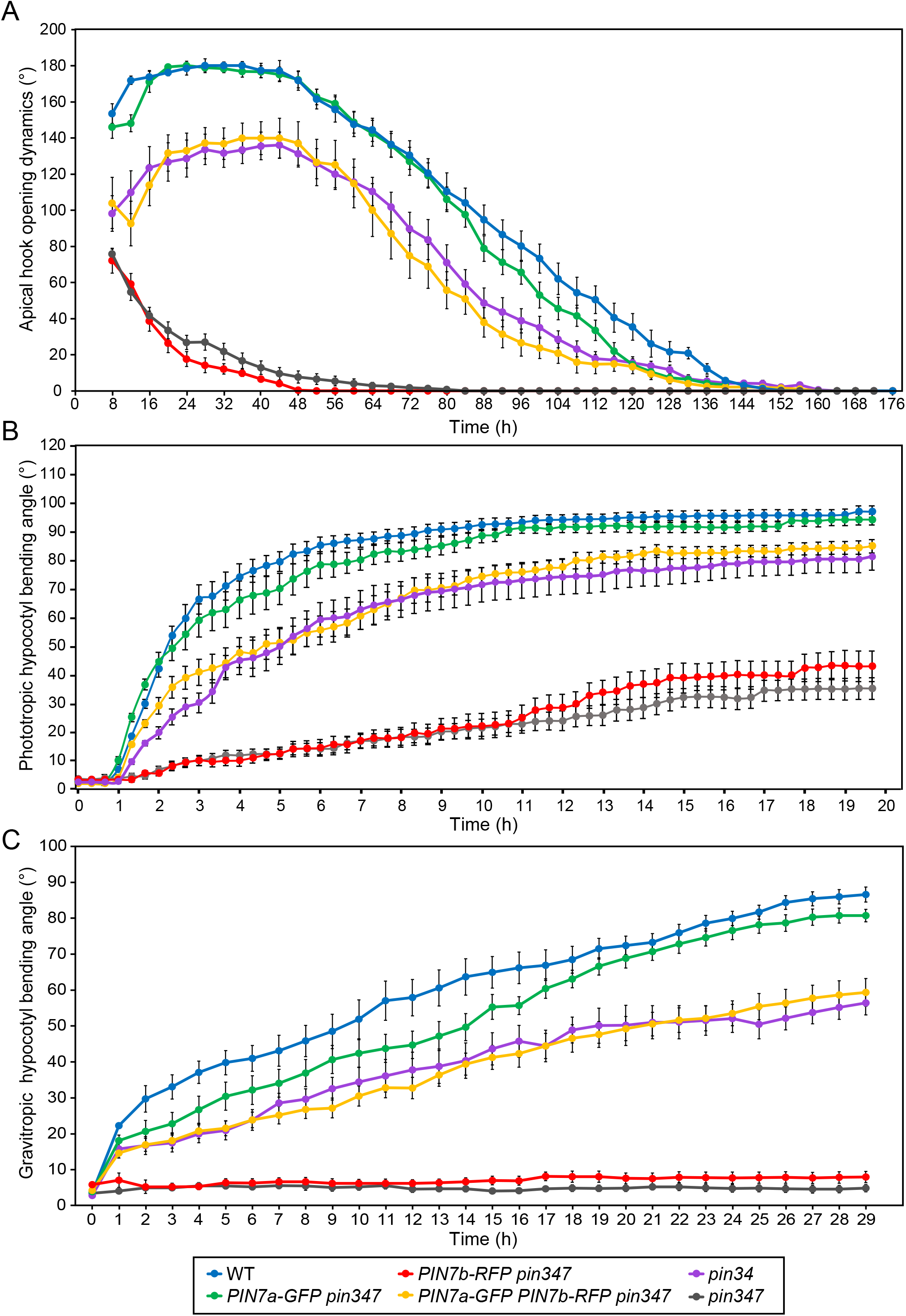
*PIN7a-GFP* and *PIN7b-RFP* cDNAs expressed in the *pin347* knock-out mutant regulate apical hook development and hypocotyl bending in a mutually antagonistic manner. **(A)** to **(C)** A temporal dynamics of etiolated *pin347* seedlings carrying both *PIN7a-*GFP and *PIN7b-RFP* cDNA transgenes during **(A)** apical hook development, and **(B)** hypocotyl phototropic and **(C)** gravitropic bending. Data are means ± S. E. For each data point, n = 15.

Apical hook development is a complex process that involves several bending steps (Zadnikova et al., 2010), and the splicing reporter suggests a slightly different expression pattern of *PIN7a* and *b* isoforms during apical hook development (Figure 2D). The hypocotyl photo- and gravitropic response includes only a single bending (Rakusová et al., 2011, 2016), and the reporter expression appears to be uniform in the respective tissues (Figure 2C). It can thus provide a hint to whether one can account for the antagonistic behavior of both isoforms to differential expression or to another mechanism, likely occurring at the cellular level. Similar to apical hook development (Figure 3A), introducing the *PIN7b-RFP* cDNA had a marginal effect on the *pin347* phenotype, while the expression of *PIN7a-GFP* lead to faster bending than that observed for the *pin34* mutant. Finally, the presence of both *PIN7a-GFP* and *PIN7b-RFP* cDNAs in *pin347* was reminiscent of the *pin34* phenotype in both phototropic and gravity assays (Figures 3B and 3C). We therefore conclude that the shared activity of both PIN7 isoforms, located probably in the same group of cells, is required for proper apical hook formation and auxin-mediated tropic responses.

### PIN7a, but not PIN7b, is required for proper formation of auxin maxima

To further investigate the functional impact of AS on PIN7, we validated the formation of the downstream auxin response maxima in apical hooks by crossing the *DR5v2:GFP* transcriptional auxin reporter (Liao et al., 2015) into lines where the phenotypic changes have been temporally monitored (see Figure 3). The reporter expression pattern in the forming apical hooks of *pin34, PIN7a-GFP pin347* and *PIN7a-GFP PIN7b-RFP pin347* resembled the situation in the wild type (Zadnikova et al., 2016), whereas the *DR5v2:GFP* distribution in the *PIN7b-RFP pin347* line was indistinguishable from that of *pin347* (Figure 4A). These results underline that the PIN7b isoform alone is unable to establish the morphogenic auxin gradients and that the observed developmental changes occur in line with previously described mechanisms (Friml et al., 2002; Willige et al., 2013; Zadnikova et al., 2010, 2016).

**Figure 4.**
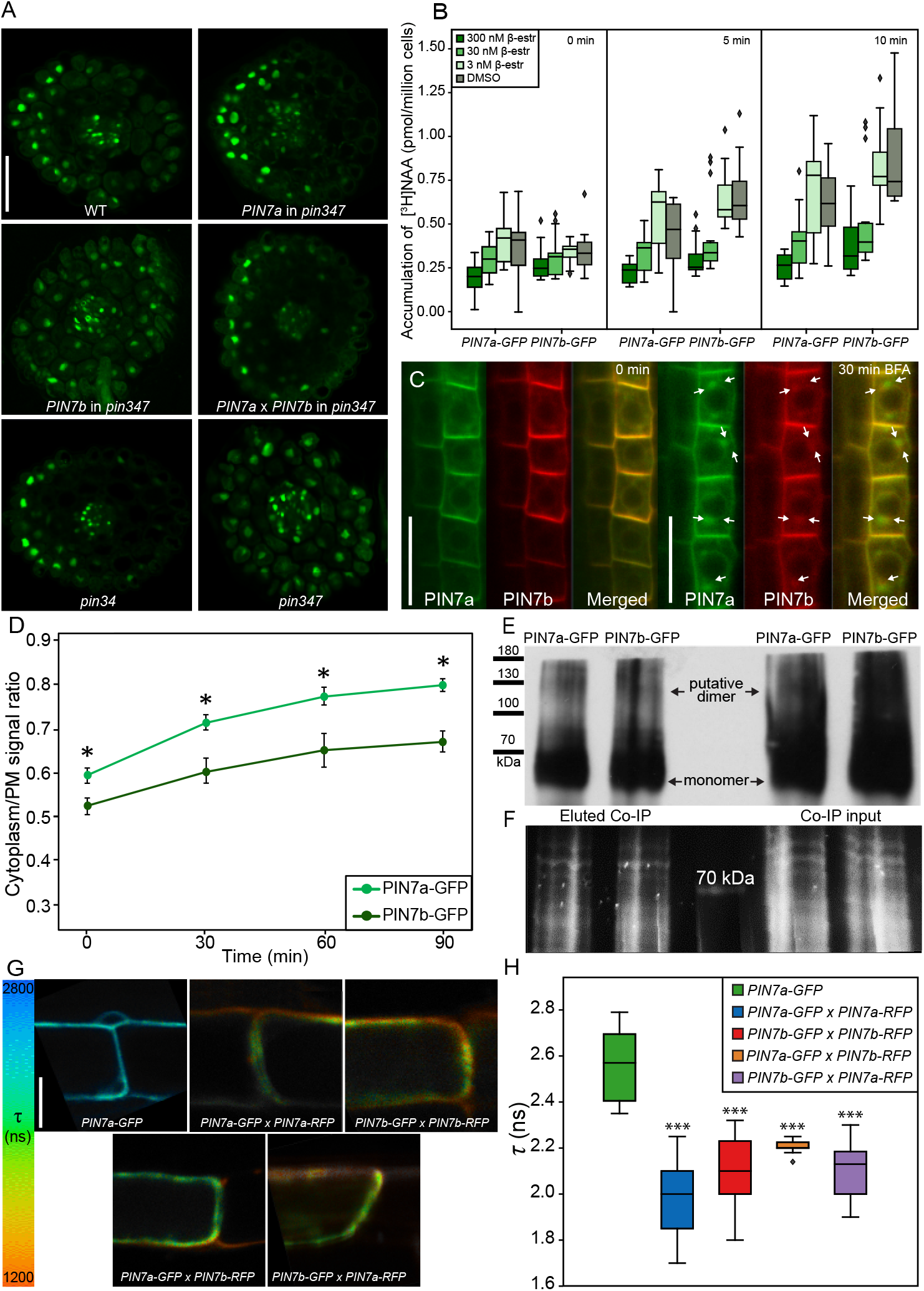
Functional analysis of the PIN7a-GFP and PIN7b-RFP interaction. **(A)** Expression of *PIN7a* but not *PIN7b* cDNA gives rise to the auxin reporter *DR5v2::GFP* gradient at the concave side of the apical hooks of the *pin347* seedlings. **(B)** Comparable [^3^H]NAA accumulation kinetics in tobacco BY-2 cells following induction of the *G10-90::XVE>>AtPIN7a-GFP* and *AtPIN7b-GFP* transgenes with decreasing doses of β-estradiol. **(C)** The effect of 50 µM brefeldin A (BFA) on the intracellular distribution of PIN7a-GFP and PIN7b-RFP. Arrows mark the incomplete co-localization of PIN7a and PIN7b in the BFA bodies. **(D)** A temporal dynamics of the BFA-mediated aggregation of PIN7a-GFP and PIN7b-GFP inside cells. The values were determined as a ratio of fluorescence intensities between the cytoplasm and PM. **(E)** and **(F)** An immunoblot (using anti-GFP antibodies) from the native-PAGE gel showing that both PIN7 isoforms can form multimeric complexes. The gel was loaded with the immunoprecipitation eluate (left) and a crude protein extract (right) from the *Arabidopsis* seedlings harboring the *G10-90::XVE>>PIN7a-GFP* and *PIN7b-GFP* constructs, induced with 5 µM β-estradiol (**[E]**, loading control **[F]**). **(G)** and **(H)** PIN7a and PIN7b can form homo-and heteromers. GFP fluorescence lifetime (*τ*) determined from the tagged PIN7a and PIN7b proteins and compared to the *τ* of the PIN7a-GFP alone, presented as a representative image heat map (**[G]** quantified on **[H]**). Proteins were expressed under the control of the *G10-90::XVE* promoter in the *Arabidopsis* root tips and induced with 5 µM β-estradiol. On the box plots, the middle line corresponds to median, the box corresponds to the 25% and 75% quantiles, the whiskers represent the minima and maxima. On **(F)**, the asterisks indicate a difference between the respective protein pair and the PIN7a-GFP control (\**P* < 0.05, \*\*\**P* < 0.001 by two-way ANOVA). On **(D)**, data are means ± S. E. For each line, on **(A)** and **(B)**, n ≥ 9, on **(C), (D), (G)** and **(H)**, n ≥ 12. Bars, 50 µm on **(A)**, 20 µm on **(C)**, and 10 µm on G.

### PIN7a and PIN7b transport auxin with comparable rates in tobacco cells

Because PIN7a and PIN7b showed differential ability to generate auxin maxima *in planta*, we asked whether both protein isoforms indeed function as auxin transporters. We expressed the *PIN7a* and *b* cDNA variants tagged with GFP under the control of the β-estradiol-driven promoter in the tobacco BY-2 tissue culture system (Petrasek et al., 2006; Müller et al., 2019). We confirmed that PIN7a (Petrasek et al., 2006), as well as the PIN7b isoform, are functional auxin transporters. To this end, we also analyzed the quantitative aspects of PIN7-mediated transport, using multiple concentrations of β-estradiol. Following induction of both transgenes, we observed a comparable decrease of radioactive-labeled auxin accumulation inside the BY-2 cells for both lines in the β-estradiol dose range used (Figure 4B), consistent with the similar PIN7a and PIN7b protein levels (Supplemental Figures 3A– 3C). The time course of the auxin accumulation drop appeared to be similar for both constructs as well (Figure 4B). Hence, *PIN7a* and *PIN7b* code for true auxin exporters, and they seem to transport auxin in tobacco cell cultures at similar rates.

### PIN7a and PIN7b differ in protein subcellular dynamics

Polarity and dynamic intracellular trafficking are essential functional attributes of PINs (Adamowski and Friml, 2015). However, the PIN7 isoforms did not prominently differ from each other in terms of polarity or general subcellular localization in the root tip at a given resolution limit (Figure 4C). The anterograde trafficking of proteins towards PM can be effectively blocked by the fungal toxin brefeldin A (BFA). It leads to internal accumulation of the membrane-bound PINs into characteristic BFA bodies (Geldner et al., 2001; Kleine-Vehn et al., 2010). During time-lapse imaging, we observed that while PIN7a-GFP accumulated readily in these intracellular aggregates, the PIN7b-RFP BFA bodies were less pronounced, co-localizing with PIN7a-GFP incompletely (Figure 4C). To exclude the effects of diverse fluorescent tags, we compared the response of PIN7a-GFP to the PIN7b-GFP cDNA lines. The BFA mediated aggregation of PIN7b-GFP indeed showed a moderate delay, compared with PIN7a-GFP (Figure 4D). It suggests that the PIN7 isoforms differ in the speed of their intracellular trafficking pathways or delivery to the PM, and the choice of the tag does not appear to interfere significantly with the subcellular dynamics of PIN7.

PIN polarity does not seem to strictly require the cytoskeleton (Glanc et al., 2019), but subcellular PIN trafficking has been proposed to be mediated by two distinct pathways (Geldner et al., 2001; Glanc et al., 2019). The first depends on actin filaments (cytochalasin D-sensitive) and occurs in most of the root meristem cells. The second (oryzalin-sensitive) utilizes microtubules and is linked with cytokinesis. Drugs that depolymerize actin filaments (cytochalasin D) and tubulin (oryzalin) (Geldner et al., 2001; Kleine-Vehn et al., 2008) showed only little effect on the intracellular localization of both PIN7 isoforms when applied alone (Supplemental Figures 3D and 3E). Pretreatment with cytochalasin D prevented the formation of the BFA bodies (Geldner et al., 2001) containing both PIN7 isoforms (Supplemental Figure 3F). Yet, when we applied oryzalin prior to the BFA treatment, the BFA compartments containing PIN7a-GFP and PIN7b-RFP associated in only very weakly co-localizing structures (Supplemental Figure 3G). Next, we tested how the cytoskeleton is involved in the trafficking of both PIN7 isoforms from the BFA bodies to the PM by washing out BFA with cytochalasin D or oryzalin (Geldner et al., 2001). In the presence of cytochalasin D, PIN7a-GFP largely persisted inside the BFA bodies, while the PIN7b-RFP signal was almost absent in these aggregates (Supplemental Figure 3H). We generally did not see any difference between both isoforms when BFA was washed out with oryzalin (Supplemental Figure 3I). These data thereby suggest that both PIN7 isoforms use vesicle trafficking pathways assisted by a common cytoskeletal scaffold. These pathways differ in their dynamics and the endomembrane components involved and are consistent with previous findings that individual PINs can utilize multiple PM delivery routes (Kleine-Vehn et al., 2008; Boutté et al., 2006).

### PIN7a and PIN7b do not differ in tropic stimuli-mediated polarity change or dimer formation

Auxin transporters of the PIN3 clade change their polar localization on the PM by the reaction to various environmental cues, in particular by switching the light or gravity vectors (Friml et al., 2002; Ding et al., 2011; Rakusová et al., 2011). PIN3 relocation in response to gravity in columella root cells is seen in as little as 2 min, while the relocation of PIN7-GFP requires approximately 30 min to be detected (Kleine-Vehn et al., 2010; Friml et al., 2002; Grones et al., 2018; Pernisova et al., 2016). We examined plants harboring both *PIN7a-GFP* and *PIN7b-RFP* cDNAs under short and long gravitropic stimuli. We did not find any difference in relocation dynamics between both isoforms in these experiments (Supplemental Figures 4A–4B). We also observed no difference in the polarity change between PIN7a-GFP and PIN7b-RFP in hypocotyl gravitropic (Rakusová et al., 2011, 2016) and phototropic (Ding et al., 2011) bending assays (Supplemental Figure 4C; due to limited transparency of hypocotyls, we used lines expressing the cDNAs under control of a stronger *SCR* promoter (Rakusová et al., 2011)). The different subcellular pathways driving both PIN7a-GFP and PIN7b-RFP cargos are thereby not connected with their ability to change polarity in response to tropic stimuli.

AS frequently changes the protein function by altering their ability to multimerize (Kelemen et al., 2013), and PIN proteins have recently been found to associate in higher-order complexes (Teale et al., 2020; Abas et al., 2021). To independently corroborate these findings, we used crude extracts from the *Arabidopsis* seedlings and the BY-2 cells, expressing the respective cDNAs, followed by non-denaturing protein electrophoresis and western blotting (Leitner et al., 2012). Probing with anti-GFP antibody indeed revealed a faint signal at the predicted size of PIN7 dimers in both tissue sources (Figures 4E and 4F; Supplemental Figures 4D and 4E). We further validated these findings with Förster resonance energy transfer by fluorescence lifetime imaging (FRET-FLIM), analyzing the cDNAs expressed stably in *Arabidopsis* seedlings and transiently in the tobacco leaves (Figures 4G and 4H; Supplemental Figures 4F and 4G). Here, we also examined the combinations of the GFP and RFP-tagged *PIN7a* and *PIN7b* cDNAs to determine their pairing preference. Shortening of the GFP fluorescence lifetime was seen regardless of the isoform interaction tested. It therefore hints at the possibility that the PIN7 splice isoforms can mutually associate into dimers (or other higher order complexes), but it seems the strength of their interaction remains comparable.

### PIN7a and PIN7b show distinct stability on PM

Numerous studies employed the fluorescence recovery after photobleaching (FRAP) analysis to investigate the dynamic turnover of various proteins, including PINs, on PM (Men et al., 2008; Martinière et al., 2012; Langowski et al., 2016; Laňková et al., 2016). We therefore bleached a region of the PM signal in the root meristem of the *PIN7a-GFP* and *PIN7b-GFP* cDNA lines and measured the FRAP in this area. Notably, PIN7a-GFP showed a slower recovery of fluorescence than PIN7b-GFP (Figure 5A); the use of either GFP or RFP tag did not markedly influence the fluorescence recovery (Supplemental Figure 5A). We also generated plants carrying a *PIN3::PIN3*Δ*-RFP* cDNA construct, which lacks the GETK motif corresponding to the four amino acids absent in PIN7b (Figures 1B and 5B). The PIN3Δ-RFP signal displayed an incomplete co-localization with the wild type PIN3-GFP variant in the BFA bodies (Supplemental Figure 5B) and faster recovery on the PM, similar to that observed for PIN7a and PIN7b (Figure 5B). It therefore appears that the motif altered by AS of PIN7 is required for the regulation of dynamics of individual isoforms within the PM.

**Figure 5.**
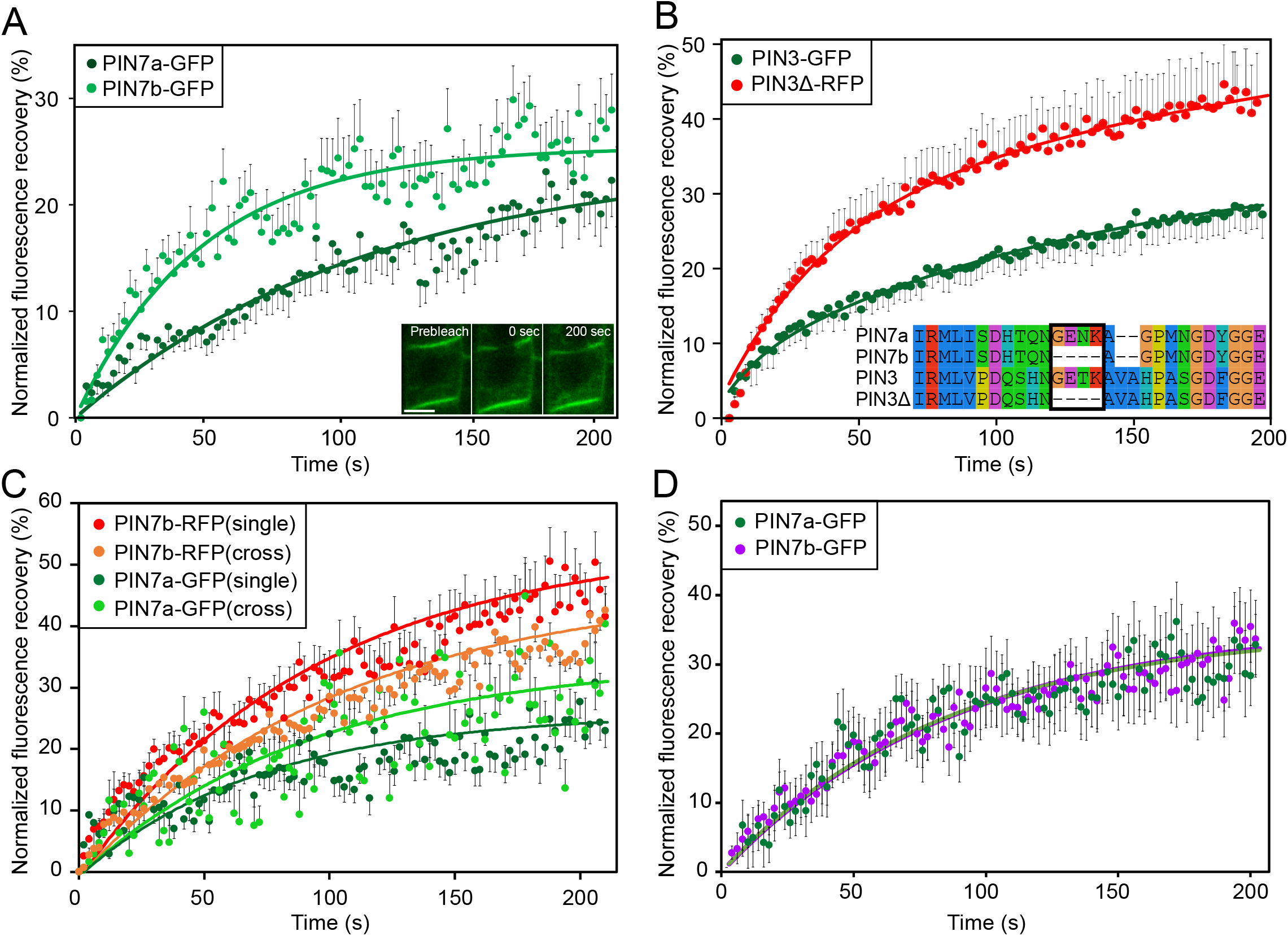
cDNA-encoded PIN7a and PIN7b mutually influence their stability within PM, as assessed by fluorescence recovery after photobleaching (FRAP). **(A)** Differential FRAP of *PIN7::PIN7a-GFP* and *PIN7::PIN7b-RFP* on PM. The example images of the bleached region (ROI) are shown on the inset. **(B)** The FRAP analysis of PIN3Δ-RFP, which lacks the 4 amino acid motif to mimic the properties of the PIN7b isoform (inset), compared to the control *PIN3-GFP* cDNA line. **(C)** The FRAP of *SCR::PIN7a-GFP* and *SCR::PIN7b-RFP* expressed in the *pin347* seedlings. The lines with a single construct are marked as “single “, the lines carrying both transgenes are marked as “cross “. **(D)** The FRAP analysis of BY-2 tobacco cell lines expressing *G10-90::XVE>>AtPIN7a-GFP* and *AtPIN7b-GFP*, following induction with 5 µM β-estradiol. Data are means ± S. E. For each data point, at least 20 ROIs were measured. The normalized data points were fit with mono-exponential curves. Bar, 5 µm on **(A)**.

The fluorophore-tagged *PIN7* cDNAs showed a relatively weak signal under natural promoter and biasing the obtained data by extensive bleaching by prolonged confocal imaging (Figure 5A). Therefore, we utilized the *SCR* promoter-driven lines, which displayed higher expression levels, but still lower intensities than the commonly used genomic DNA-derived *PIN7:PIN7-GFP* (Blilou et al., 2005) (Supplemental Figures 5C and 5D). Indeed, these lines allowed for detecting a finer difference in FRAP, as compared to those under the control of the native promoter (Figure 5C). The FRAP experiments (Figure 5A) were conducted initially on the *pin347* lines, which expressed the corresponding transgenes separately. Therefore, we compared the FRAP values of the single PIN7a-GFP and PIN7b-RFP with those carrying these constructs simultaneously in the *pin347* genetic background. Strikingly, the co-expression of PIN7a-GFP decreased the FRAP of PIN7b-RFP and, *vice versa*, the presence of PIN7b-RFP increased the FRAP of PIN7a-GFP (Figure 5C). This shows that the PIN7 isoforms mutually influence each other stability within PM, presumably by the association in a molecular complex.

In addition, both PIN7a and PIN7b did not markedly differ in their capability to transport auxin in the BY-2 cells (Figure 4B and Supplemental Figure 3A). We were unable to detect homologous AS events in plant lineages related to tobacco (see Figure 1D). We tested FRAP of the *Arabidopsis* PIN7a- and PIN7b-GFP on PM in BY-2 cells. Interestingly, the recovery of both isoforms showed a virtually identical course (Figure 5D). Altogether, these observations support the scheme that both isoforms physically interact and mutually influence each other stability within PM, which leads to a differential impact on the plant phenotype.

## DISCUSSION

We provide evidence that the evolutionary conserved AS event increases the diversity among the PIN auxin transporters in *Arabidopsis thaliana*. Focusing on PIN7, we show that both PIN7a and PIN7b isoforms (differing by presence of a 4-amino acid stretch) are required together for the correct apical hook formation and tropic responses. A similar, genetically evidenced cooperative modes of action of splice isoforms has already been proposed during *Arabidopsis* development (Szakonyi and Duque, 2018). The seed dormancy regulator *DELAY OF GERMINATION 1* (*DOG1*) is processed into five mRNAs. Only the expression of two or more DOG1 cDNAs under the native promoter rescues the *dog1* phenotype by synergistic stabilization of the protein by its multimerization (Nakabayashi et al., 2015). The transcriptional factor HYH (HY5 HOMOLOG) possesses an isoform, which lacks a domain required for proteasomal degradation, which leads to its increased stability and probably works as a semi-dominant splice variant (Szakonyi and Duque, 2018; Sibout et al., 2006); this can probably also be the case of the AS of *MONOPTEROS* (Cucinotta et al., 2020). Together with a partly similar mechanism described for the temperature-mediated regulation of flowering (Posé et al., 2013; Lee et al., 2013), the example of AS of PIN7 is among rarely described instances where the mutually antagonistic effects of two splice isoforms are observed and can be placed in developmental context.

Based on the current understanding of the PIN-mediated auxin transport (Adamowski and Friml, 2015), we validated the most plausible hypotheses to provide a mechanistic view on the mutual functionality of PIN7a and PIN7b. This AS event neither affects the ability of PIN7 carriers to transport auxin *per se* nor influences their ability to relocate after the tropic stimulus. Although the expression of PIN7b is pronounced at the concave side of the apical hook, we did not see any remarkable expression changes of the P7A_1_G/P7BR dual reporter during hypocotyl bending assays (Figure 2C and Supplemental Figure 4C). Instead, our data favor the scenario that PIN7a and PIN7b functionally interact at the protein level. Along with recent findings (Teale et al., 2020; Abas et al., 2021), we confirm with orthogonal experimental tools that PIN7 isoforms closely associate. Moreover, the combined FRAP analysis shows that PIN7a and PIN7b influence each other stability within PM. It has previously been suggested that PINs are complexed inside stable integral PM clusters (Langowski et al., 2016; Feraru et al., 2011; Kleine-Vehn et al., 2011). The concept that PIN7a, detained in these microdomains, transports auxin, but dynamic PIN7b does not, can also be in accord with the diverse subcellular pathways delivering the PIN7 isoforms to PM. Hence, PIN7b probably binds PIN7a and impedes the polar auxin flow provided by PIN7a in these membrane domains to fine-tune auxin developmental responses (Supplemental Figure 5E).

In general, AS modifies protein properties variably. It can affect protein subcellular localization, ligand binding affinity, enzymatic or transporting activities, protein stability, or the presence of covalent post-translational modifications (Stamm et al., 2005; Kelemen et al., 2013). Different covalent modifications alter the subcellular trafficking of most PINs. Phosphorylation sites on serine, threonine or tyrosine residues of various PINs have been identified; their phosphorylation status also changes PIN-mediated tropic responses (Zwiewka et al., 2019; Barbosa et al., 2018; Rademacher and Offringa, 2012). Also PIN ubiquitination (on lysines) and controlled proteolytic degradation act in auxin-mediated processes (Leitner et al., 2012). However, none of the candidate residues required for these modifications is present in the vicinity of the amino acid motif changed by AS. This region, present inside the PIN long hydrophilic loop, shows low amino acid conservation (as seen particularly on the *PIN4* AS event), and it is perhaps intrinsically disordered. Therefore, one can speculate whether just the length of this motif can be critical for functional interaction among internal structural domains (Buljan et al., 2013). These structural differences may modulate the affinity of PINs to bind the factors required for the entry and presence in the PM microdomains, and thus participate in the PIN subcellular dynamics and directional auxin transport.

## EXPERIMENTAL PROCEDURES

### Plant material and plant growth conditions

All plant material, except tobacco cell cultures, was in the *Arabidopsis thaliana* (L.) Heynh., Col-0 ecotype. These mutant and transgenic lines were described previously: *PIN3::PIN3-GFP* (Zadnikova et al., 2010), *PIN4::PIN4-GFP, PIN7::PIN7-GFP* (Blilou et al., 2005), *pin3-3 pin4-101 pin7-102* (*pin347*), *pin3-3 pin4-101* (*pin34*) (Willige et al., 2013), *pin7-2* (Friml et al., 2003).

For the *in vitro* cultivation, the seeds were surface-sterilized for 5 h with chlorine gas, plated on 0.5× Murashige & Skoog medium with 1 % sucrose, and then stratified for 2 d at 4°C in darkness. Unless indicated otherwise, the seedlings were grown on vertically oriented plates for 4-6 days under 16 h: 8 h photoperiod, 22: 18°C, light: dark.

The following chemicals were used for treatments: brefeldin A (BFA), cytochalasin D (CytD), oryzalin (Ory), β-estradiol, 2,4-dichlorophenoxyacetic acid (2,4-D), all from Sigma (Sigma-Aldrich, St. Louis, MO, USA). Radioactively labeled auxin accumulation assays were performed with [^3^H]NAA (naphthalene-1-acetic acid; 20 Ci.mmol^-1^; American Radiolabeled Chemicals, St. Louis, MO, USA).

### DNA manipulations and transgenic work

The genomic *PIN7::PIN7-RFP* construct was made by replacing the GFP coding sequence in the original *PIN7::PIN7-GFP* construct (Blilou et al., 2005). For creating the *PIN7(4)*::*PIN7(4)a-GFP* and *PIN7(4)*::*PIN7(4)b-RFP* cDNA constructs, the respective cDNAs were cloned into the pDONR221 P5-P2 entry vector (Invitrogen, Life Technologies, Carlsbad, CA, USA) by the Gateway BP reaction (Invitrogen). For placing the fluorophore tag, the *Xba*I restriction site was introduced at the 1350 bp (for *PIN7*) or 1341 bp (for *PIN4*) behind the start codon of the cDNA coding region, in the same amino acid sequence context as in the published *PIN4* and *PIN7-GFP* constructs (Blilou et al., 2005). In parallel, the *PIN7* (or *PIN4*) promoters were inserted into the pH7WG Gateway vector (Department of Plant Systems Biology, Ghent University, Belgium (Karimi et al., 2002)) by the Gibson Assembly kit (New England Biolabs, Ipswich, MA, USA). The tagged *PIN7* and *PIN4* cDNA entry clones were then recombined with the modified pH7WG destination vector by the Gateway LR reaction (Invitrogen). For the cloning of the *PIN7* splicing dual fluorescent reporter, the entry vectors carrying the genomic sequence of *PIN7-GFP* and *PIN7-RFP*, respectively, were modified by inverse PCR. The resulting constructs were then recombined by the Gateway LR reaction with the *PIN7* promoter containing the pH7WG destination vector. The *SCR::PIN7a-GFP* and *SCR::PIN7b-RFP* constructs were obtained by recombination of the *SCR* promoter in pDONR221 P1-P5r and *PIN7* cDNA entry clones with the pH7WG destination vector by the Multisite Gateway LR reaction. *PIN3*Δ*-RFP* cDNA was custom synthesized (Gen9, Ginkgo Bioworks, Boston, MA, USA), cloned into pDONR221 P5-P2 vector and together with the *PIN3* promoter pDONR221 P1-P5r construct recombined into the pH7WG vector with Multisite Gateway. The validated binary constructs were transformed into *Arabidopsis* by floral dipping. For making the estradiol-inducible *G10-90::XVE>>PIN7(4)a-GFP, PIN7(4)b-GFP, PIN7a-RFP* and *PIN7b-RFP* constructs, tagged *PIN7* or *PIN4* cDNA entry clones were recombined with the pMDC7 destination vector (Curtis and Grossniklaus, 2003) by the Gateway LR reaction. Unless stated otherwise, the constructs present in the study were cloned under their natural *PIN4* or *PIN7* promoter. Primers used in this work are listed in Supplemental Table 1.

For the generation of the stable transgenic lines carrying the cDNA constructs, at least eight independent descendant populations of primary transformants were preselected for the presence of the fluorescent signal. The functional validity of the cDNA constructs was verified in the phototropic bending test, where all candidate lines matched the presented results. Selected lines were used for further phenotypic analysis.

### Plant phenotype analysis

The dynamic seedling development was tracked in the custom made dynamic morphogenesis observation chamber equipped with blue and white LED unilateral light sources, infra-red LED back light and a camera for imaging in the infra-red spectra, controlled by the Raspberry Pi3B microcomputer (Raspberry Pi foundation, Cambridge, UK). For the hypocotyl bending assays, the plated seeds were first illuminated for 6 h with white light. The plates were then transferred to the observation chamber for 3-4 d. For the hypocotyl phototropic bending experiments (Friml et al., 2002), the dark-grown seedlings were afterward illuminated for 20 h with unilateral white light and imaged every 20 minutes. For hypocotyl gravitropic bending experiments (Rakusová et al., 2011), the dark-grown seedlings were rotated by 90° clockwise and imaged for 30 h every 60 min. For tracking apical hook development, the seeds were first illuminated for 6 h with white light. They were then transferred to the observation chamber and their development recorded every 4 h for a total 150 - 200 h in infrared spectra. Germination time was set as time 0, when first traces of the main root were observed, individually for each seedling analyzed. Apical hook was determined as an angle between the immobile (non-bending) part of hypocotyl and the distal edge of cotyledons (Zadnikova et al., 2010), using the ImageJ software (Rueden et al., 2017). At least 15 seedlings were analyzed for each line. Each experiment was done at least three times.

For protoxylem defects analysis, 5-d old light-grown seedlings were cleared and analyzed as described previously (Bishopp et al., 2011). For examining lateral root density, 8-d old light-grown seedlings were cleared and observed with a DIC microscope (Dubrovsky et al., 2009). Lateral root density was calculated by dividing the total number of lateral roots and lateral root primordia to the length of the main root as described (Dubrovsky et al., 2009). Vertical Growth Index (VGI), defined as a ratio between main root ordinate and main root length, was quantified on 5-d old seedlings as published previously (Grabov et al., 2005). For measuring the Gravitropic Set-point Angle (GSA), plates with 14-d old light-grown seedlings were scanned and the angles between the vertical axis and five innermost 0.5 mm parts of lateral root were determined as described previously (Roychoudhry et al., 2017). In all cases, 12-20 seedlings were analyzed for each line. Each experiment was done at least three times.

For decapitation experiments, 4-week short-day grown plants were moved into long-day conditions to induce flowering. After the primary bolt reached 10-15 cm, the plant was decapitated. The number of rosette branches was recorded at 7, 10 and 14-d after decapitation (Waldie and Leyser, 2018; Greb et al., 2003). Rosette size was inspected in 18-d light-grown plants prior to documenting. 10 plants were analyzed for each line, the experiment was done three times.

### Microscopy

Bright-field microscopy (differential interference contrast, DIC) was conducted on the Olympus BX61 instrument (Olympus, Shinjuku, Tokyo, Japan) equipped with a DP50 camera (Olympus). Routine confocal microscopy was performed on inverted Zeiss Axio Observer.Z1 containing the standard confocal LSM880 and Airyscan modules with 20x/0.8 DIC M27 air, 40x/1.2 W Kor FCS M27 air and 63x/1.40 Oil DIC M27 objectives (Carl Zeiss AG, Jena, Germany). Gravity-induced polarity change experiments were carried out on Zeiss Axio Observer.Z1 with vertically oriented sample position and the 40x/0.75 glycerol objective.

To observe the light-induced polarity change of PIN7a-GFP and PIN7b-RFP, 4-d dark-grown seedlings were irradiated for 4 h with unilateral white light and then imaged with Zeiss Axio Observer.Z1 LSM880 with a vertically oriented sample position, as described (Willige et al., 2013). For analyzing gravity-induced polarity change, 4-d dark-grown seedlings were reoriented by 90° clockwise and imaged 6 and 24 h after rotation, as described (Rakusová et al., 2011).

For BFA treatments, 5-d light-grown seedlings were transferred to liquid 0.5× MS media containing 50 μM BFA. The membrane/cytosol ratio was determined with ImageJ, it was defined as the mean membrane signal intensity divided by the mean fluorescence in the cytosol. For cytoskeleton depolymerizing drug treatments (Geldner et al., 2001), 5-d light-grown seedlings were transferred to liquid 0.5× MS media supplemented with 20 μM of cytochalasin D or 20 μM oryzalin. The co-treatments were done by the direct addition of BFA. For the BFA removal, prior to the addition of the cytoskeleton depolymerizing compounds, seedlings were twice washed out with fresh media and then transferred to that supplemented with the respective cytoskeleton-depolymerizing drug.

For imaging of the expression of *DR5v2:GFP* in the apical hook, the GFP signal in the 4-days old etiolated seedlings was fixed in 4% paraformaldehyde (PFA, Sigma) overnight. Then, the tissue was hand-sectioned with a razor blade and observed.

### Fluorescent recovery after photobleaching (FRAP)

For the FRAP experiments, Zeiss Axio Observer.Z1 equipped with the LSM880 confocal and Airyscan modules and the 40x/1.2 W Kor FCS M27 air objective was used. A rectangular region of interest (ROI) of 40 × 20 pixels was selected on the basal or apical PM of cells inside the vascular cylinder in the primary root meristematic area of 5-d light-growth seedlings. ROI was bleached with the maximum 488 nm laser intensity and fluorescence recovery was documented every 1 or 2 sec for a total 200 sec. Recovery time lapses were analyzed with ImageJ. The Slices Alignment plugin (Tseng et al., 2012) for ImageJ was used for the elimination of cell movement caused by root growth. In parallel, another rectangular ROI (100 × 20 pixels) on the non-bleached cell was selected as a reference. To compensate for the fluorophore bleaching during the recovery period, the data were normalized using equation (Laňková et al., 2016)

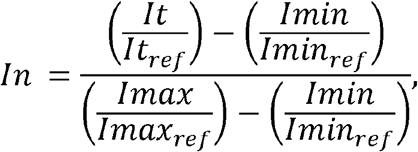

where *In* is normalized fluorescence intensity, *It* is the intensity at the specific time point, *Imax* is the intensity after the initial bleaching, *Imax* is the intensity before initial bleaching. *It*_*ref*_, *Imax*_*ref*_ and *Imin*_*ref*_ represent the same values for the reference ROI. At least 3 membranes in 4-5 root tips were analyzed in each experiment. Single-phase exponential fitting was applied to the normalized FRAP data (SigmaPlot, Systat Software, Chicago, IL, USA) as described (Laňková et al., 2016; Sprague and McNally, 2005). Recovery half-time is defined as a time required for the fluorescence recovery to reach half of the steady-state fluorescence intensity (Sprague and McNally, 2005; Soumpasis, 1983).

### FLIM-FRET analysis

For assessing the pairing specificity in *Arabidopsis*, the lines harboring all 4 combinations of *G10-90::XVE>>PIN7a/b-GFP* (donors) and *PIN7a/b-RFP* (acceptors) were generated. Before imaging, 4-d light-growth seedlings were transferred for two days to the solid ½ MS media supplemented with 5 μM β-estradiol. The transient expression in tobacco leaves was performed as earlier published (Horák et al., 2008), omitting the β-estradiol induction to keep the transgene expression low (Bashandy et al., 2015). The abaxial epidermis of tobacco leaves was observed 3 days after infiltration.

The procedure of the FRET-FLIM assay was done as previously described (Krejčí et al., 2015). Imaging was performed on inverted Zeiss LSM 780 AxioObserver.Z1 equipped with the M27 Plan-Apochromat 20x air objective, external In-Tune laser (<3 nm width, pulsed, 40 MHz, 1.5 mW) and the GaAsP detectors. For conducting the FLIM-FRET assays, the HPM-100-40 hybrid detector associated with the photon counting module Simple-Tau 150 (compact TCSPC system based on the SPC-150N device) was utilized, using the detector controller card DCC-100 (Becker & Hickl GmbH, Berlin, Germany). The cell membranes containing both donor and acceptor signals were excited with the wavelength 490 nm to generate intensity image and the photons were recorded with the supplied SPCM software. In the image analysis, the pixel per pixel fitting model was used for the two-component system and the one-component system was used for the donor alone (Becker & Hickl SPCImage software package). Multiexponential decay model was used for fitting and FRET efficiencies were calculated using the equation

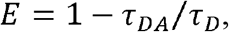

where *E* is energy transfer efficiency, *τ*_DA_ donor fluorescent lifetime in the presence of acceptor and *τ*_*D*_ donor fluorescent lifetime in the absence of acceptor. The observation of two lifetimes in the GFP fluorescence spectrum was expected: first, close to 2.6 ns, corresponds to a non-interacting fraction, while the second one, characterized by a shorter lifetime, was considered as associated with interacting molecules.

### Tobacco cell lines and auxin accumulation assays

Tobacco cell line BY-2 (*Nicotiana tabacum* L. cv. Bright Yellow 2) was cultivated as described (Müller et al., 2019). BY-2 cells were transformed with the pMDC7 constructs by co-cultivation with *Agrobacterium tumefaciens* strain GV2260 as earlier described (Petrasek et al., 2006; Müller et al., 2019). The transgene expression was maintained by cultivation on media supplemented with 40 μg.ml^-1^ hygromycin (Roche) and 100 μg.ml^-1^ cefotaxime (Sigma). Accumulation assays of radioactively labeled auxin were performed as previously published (Petrasek et al., 2006; Delbarre et al., 1996) with cells cultured for 2 days in β-estradiol or DMSO mock treatments. Presented results are from 3 biological replicates for each representative *PIN7a* and *PIN7b* line and were confirmed for each on two other independent genotypes. Independent β-estradiol inductions were done within 3-9 months after establishing the cell suspension on lines, which showed comparable levels of the expressed PIN7-GFP signal.

### Protein extraction and immunoblotting

Protein extraction from the BY-2 tobacco cell lines and the *Arabidopsis* roots was performed as described previously (Leitner et al., 2012), using the modified buffer, which contained only NP-40 as detergent. Otherwise, DTT, SDS and any other detergents were omitted from the buffers. The GFP tagged PIN7 variants were immunoprecipitated with the GFP-Trap Dynabeads (Chromotek, Planegg, Germany) according to the manufacturer ‘s description. The bead elution was done with the glycine buffer (0.2 M glycine, pH 2.5) and the eluate was neutralized with the 1/10 sample volume of 1M Tris-HCl, pH 8.5. The eluted proteins were mixed with a premade 2× Sample Buffer (Biorad, Hercules, CA, USA), omitting β-mercaptoethanol (Sigma) prior to the loading on the gel. To determine the putative dimerization of PIN7-GFP, the total membrane fractions or immunoprecipitated proteins were run on a non-denaturing polyacrylamide gel electrophoresis. Here, the Mini-PROTEAN® TGX™ Stain-Free Precast gels (Biorad) were used, devoid of SDS. The trihalo compounds present in the gel allowed for the visualization of the proteins under UV light to control the loading quality. As a marker, the PageRuler™ Prestained Protein Ladder (10 to 180 kDa, ThermoFisher) was used. For western blotting, the proteins were transferred to the nitrocellulose membrane (GE Healthcare Life Sciences, Chicago, IL, USA), blocked with 3% BSA for 2 h, and incubated overnight at 4°C with monoclonal primary anti-GFP antibody from rabbit (1:2,000) (Invitrogen), and with secondary anti-rabbit antibody (1:30,000) (Invitrogen), as described previously (Leitner et al., 2012).

### Statistics and sequence analysis

For the mean comparison of the two groups, Student ‘s *t*-test was applied. Statistical analysis of multiple groups was performed by one-way ANOVA with subsequent Tukey HSD post-hoc test. All statistical tests, including that for equal normality, were performed in R-studio IDE (R-studio, Boston, MA, USA). In the box plots, the whisker length was set as *Q* ± × IQR, where *Q* is the corresponding quartile and IQR is the interquartile range. For creating the multiple sequence alignments, the protein sequences were assembled with the Clustal Omega algorithm (Sievers et al., 2011) and graphically outlined by Mega-X, using default ClustalX color code (Kumar et al., 2018).

## ACKNOWLEDGMENT

We thank Claus Schwechheimer for the *pin34* and *pin347* seeds, Yuliia Mironova for technical assistance, Ksenia Timofeyenko and Dmitry Konovalov for help with the evolutional analysis, Konstantin Kutashev and Siarhei Dabravolski for assistance with the FRET-FLIM analysis, Huibin Han for advice with the hypocotyl imaging, Karel Müller for the initial qRT-PCR on the tobacco cell lines, and Jozef Mravec and Lindy Abas for their comments on the manuscript. This work was supported by the Czech Science Foundation (16-26428S and 19-23773S) to I. K., M. H. and K. R, and the Ministry of Education, Youth and Sports of the Czech Republic (MEYS, CZ.02.1.01/0.0/0.0/16_019/0000738) to K. R. The imaging Facilities of the Institute of Experimental Botany and CEITEC are supported by MEYS (LM2018129 - Czech BioImaging and CZ.02.1.01/0.0/0.0/16_013/0001775).

## AUTHOR CONTRIBUTIONS

K., M. H., K. Re., J. Hu., A. J., R. F., Z. V., D. R., and J. P. conducted experiments. S. S. and J. F. provided unpublished material. I. K., K. Re., J. Hu., J. P., T. B. J., M. J. F., J. He., J. F., J. P., and K. Rů. conceived the research and designed experiments. K. Rů. and I. K. wrote the manuscript. All authors read and commented on the final version of the manuscript.

### DECLARATION OF INTERESTS

The authors declare no competing interests.

## Notes

### Competing Interest Statement

The authors have declared no competing interest.

