## Supplemental Figure 1 for "Mutually opposing activity of PIN7 splicing isoforms is required for auxin-mediated tropic responses in *Arabidopsis thaliana*"

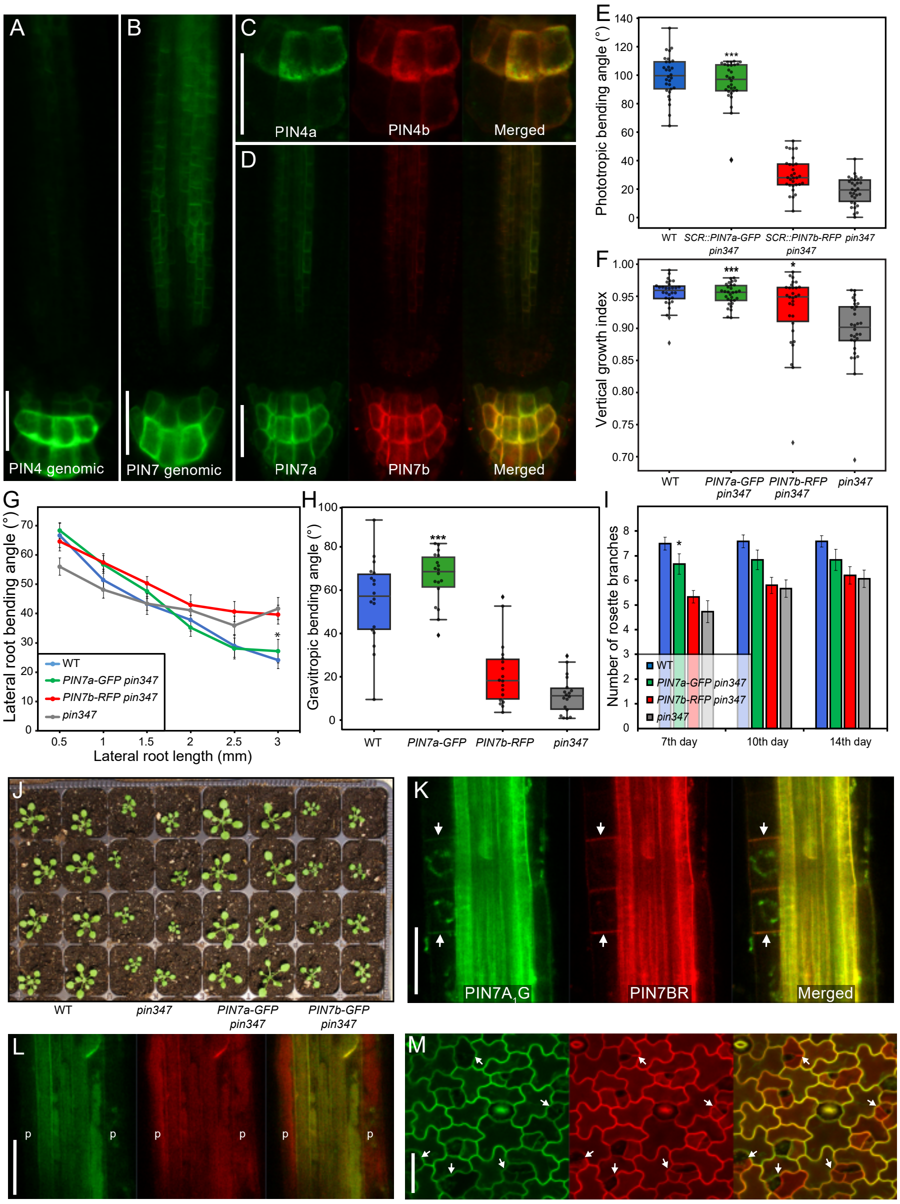


**Supplemental Figure 1.** The localization, phenotypic and expression features of the PIN7a and PIN7b splice isoforms.

**(A)** to **(D)** The expression and localization pattern of the genomic *PIN4::PIN4-GFP* **(A)** and *PIN7::PIN7-GFP* **(B)**, compared with the *PIN4::PIN4a-GFP*, *PIN4::PIN4b-RFP* **(C)**, *PIN7::PIN7a-GFP* and *PIN7::PIN7b-GFP* **(D)** cDNA constructs in the primary root tip.

**(E)** Phototropic bending of the hypocotyl of the *pin347* mutants harboring the *PIN7a-GFP* and *PIN7b-RFP* cDNAs under control of the *SCR* promoter.

**(F)** to **(J)** Additional phenotypes of the *pin347* plants carrying the *PIN7a-GFP* and *PIN7b-RFP* cDNAs. This includes tests for root gravitropism **(F)**, the gravitropic set-point angle, as a parameter of orthogravitropism of lateral roots **(G)**, hypocotyl bending angle induced by gravity **(H)**, the number of rosette branches arising after decapitation **(I)** and the rosette stage development **(J)**.

**(K)** to **(M)** Differential expression of the $P7A_1G$/P7BR reporter in the lateral root primordia **(K)**, in the pericycle cells (p) adjoining phloem in the mature primary root **(L)**, and in the stomatal lineage ground cells of the cotyledon epidermis **(M)**.

On the box plots, the middle line corresponds to median, the box to the 25% and 75% quantiles, the whiskers represent the minima and maxima. On **(G)** and **(I)**, the data are means ± S. E. Asterisks represent a difference between the respective line and the *pin347* mutant (*$P < 0.05$, ***$P < 0.001$ by two-way ANOVA). On (A–D) and (K–M), at least 9 samples were analyzed, for each line on (E–I), at least 15 samples were quantified. Bars, 25 µm on **(A)** to **(D)** and **(K)** to **(M)**.
