## Supplemental Figure 2 for "Mutually opposing activity of PIN7 splicing isoforms is required for auxin-mediated tropic responses in *Arabidopsis thaliana*"

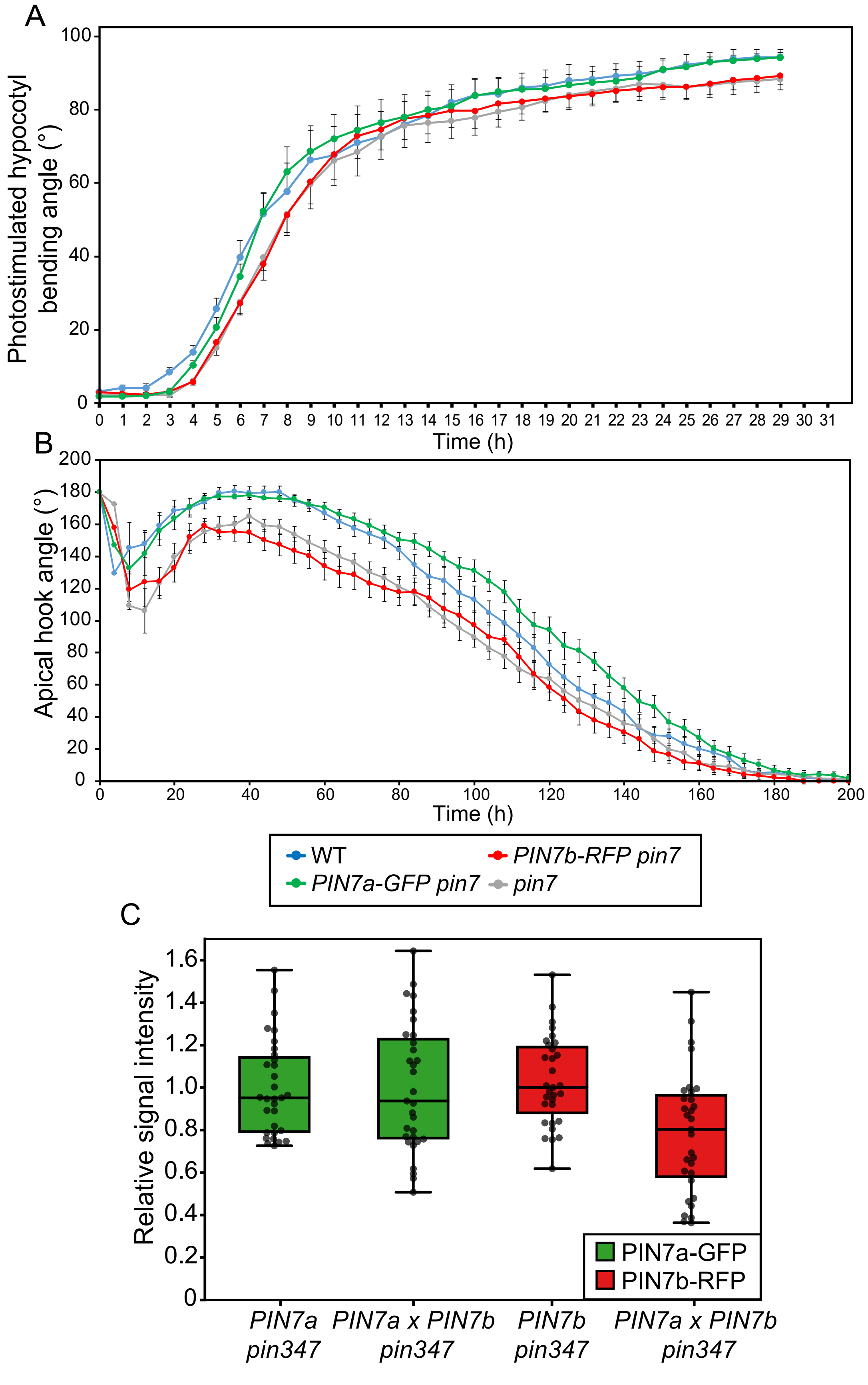


**Supplemental Figure 2.** Additional *PIN7* cDNA complementation and expression tests.

**(A)** and **(B)** A subtle difference between the wild type and *pin7-2* mutants expressing the *PIN7a-GFP* and *PIN7b-RFP* cDNAs in the temporal analysis of the phototropic bending **(A)** and during apical hook development **(B)**. The data are means ± S. E. For each line, 15 seedlings were measured.

**(C)** A comparison of the fluorescence intensities in the *pin347* mutants carrying the combinations of the *PIN7a-GFP* and *PIN7b-RFP* cDNAs. The values were not different at the given significance level ($P$ > 0.05 by Student's $t$-test, 30 cells analyzed for each line). The box corresponds to the 25% and 75% quantiles, whiskers represent the minima and maxima, dots denote single data points.
