## Supplemental Figure 3 for "Mutually opposing activity of PIN7 splicing isoforms is required for auxin-mediated tropic responses in *Arabidopsis thaliana*"

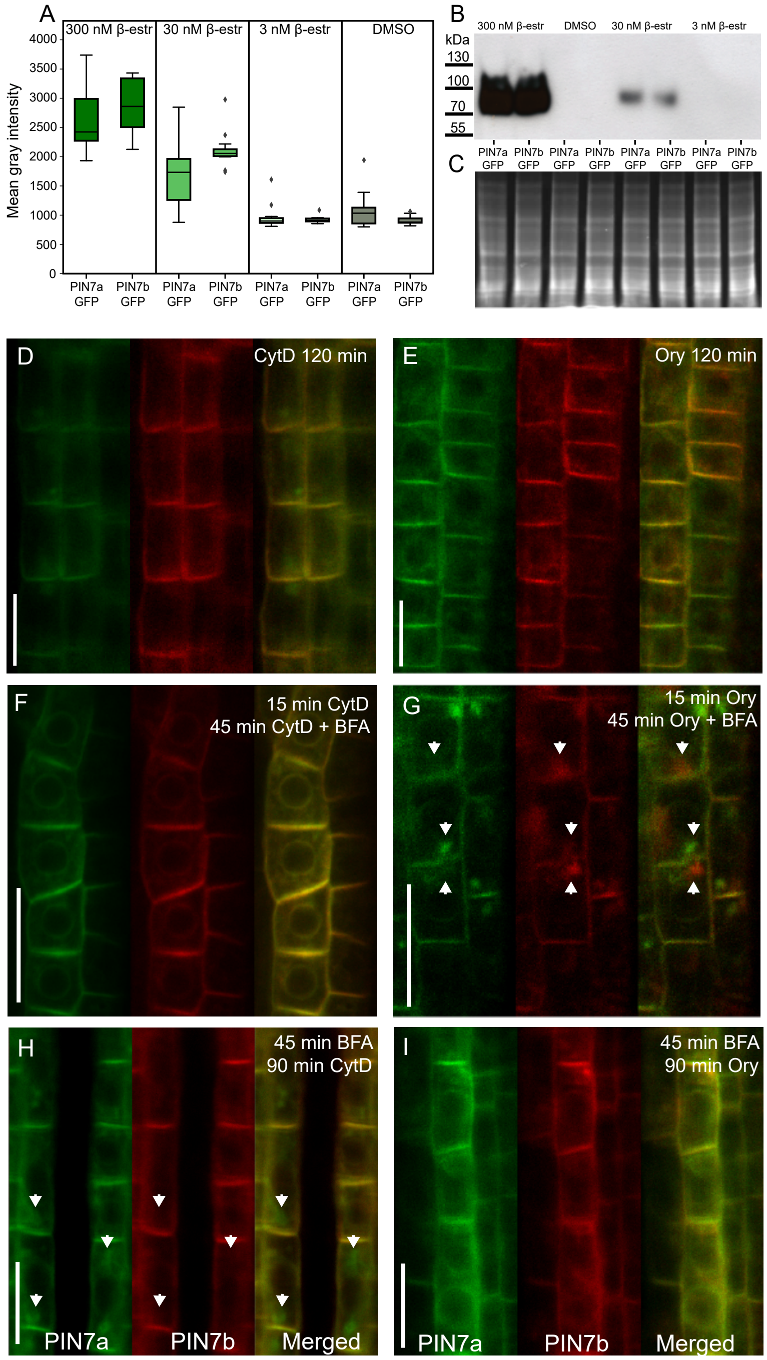


**Supplemental Figure 3.** Quantitative and qualitative properties of the PIN7a and PIN7b cDNA-based proteins expressed in the BY-2 cells and *Arabidopsis* seedlings.

**(A)** to **(C)** Quantification of the *G10-90::XVE>>AtPIN7a-GFP* and *AtPIN7b-GFP* levels in BY-2 cells, following induction with decreasing doses of β-estradiol. (A) Mean gray intensities of the total image membrane fluorescent signal. Immunoblot of total protein extracts from the transgenic BY-2 cells, detected with anti-GFP antibody (**[B]**, loading control **[C]**).

**(D)** and **(E)** Absence of remarkable effects on the PIN7a-GFP and PIN7b-RFP localization after 2-h treatments with the 20 µM actin polymerization inhibitor cytochalasin D **(D)** and with the 20 µM microtubule depolymerization drug oryzalin **(E)**.

**(F)** The fluorescence signal from PIN7a-GFP and PIN7b-RFP upon disruption of actin filaments with 20 µM cytochalasin D for 15 min, followed by the addition of 50 µM BFA for another 45 min.

**(G)** PIN7a-GFP and PIN7b-RFP aggregation (arrows) upon disruption of microtubules with 20 µM oryzalin for 15 min, followed by the addition of 50 µM BFA for another 45 min.

**(H)** Effect of cytochalasin D on the PIN7a-GFP and PIN7b-RFP exit from the BFA bodies upon 45 min pretreatment with 50 µM BFA and followed by wash-out with 20 µM cytochalasin D for another 90 min.

**(I)** Effect of oryzalin on the PIN7a-GFP and PIN7b-RFP exit from the BFA bodies upon 45 min pretreatment with 50 µM BFA and followed by wash-out with 20 µM oryzalin.

On **(A)**, the box corresponds to the 25% and 75% quantiles, whiskers represent the minima and maxima, for each line, 10 images were analyzed. On **(D)** to **(I)**, 3 images from each 4 root tips were analyzed. Bars, 10 µm.
