## Supplemental Figure 4 for "Mutually opposing activity of PIN7 splicing isoforms is required for auxin-mediated tropic responses in *Arabidopsis thaliana*"

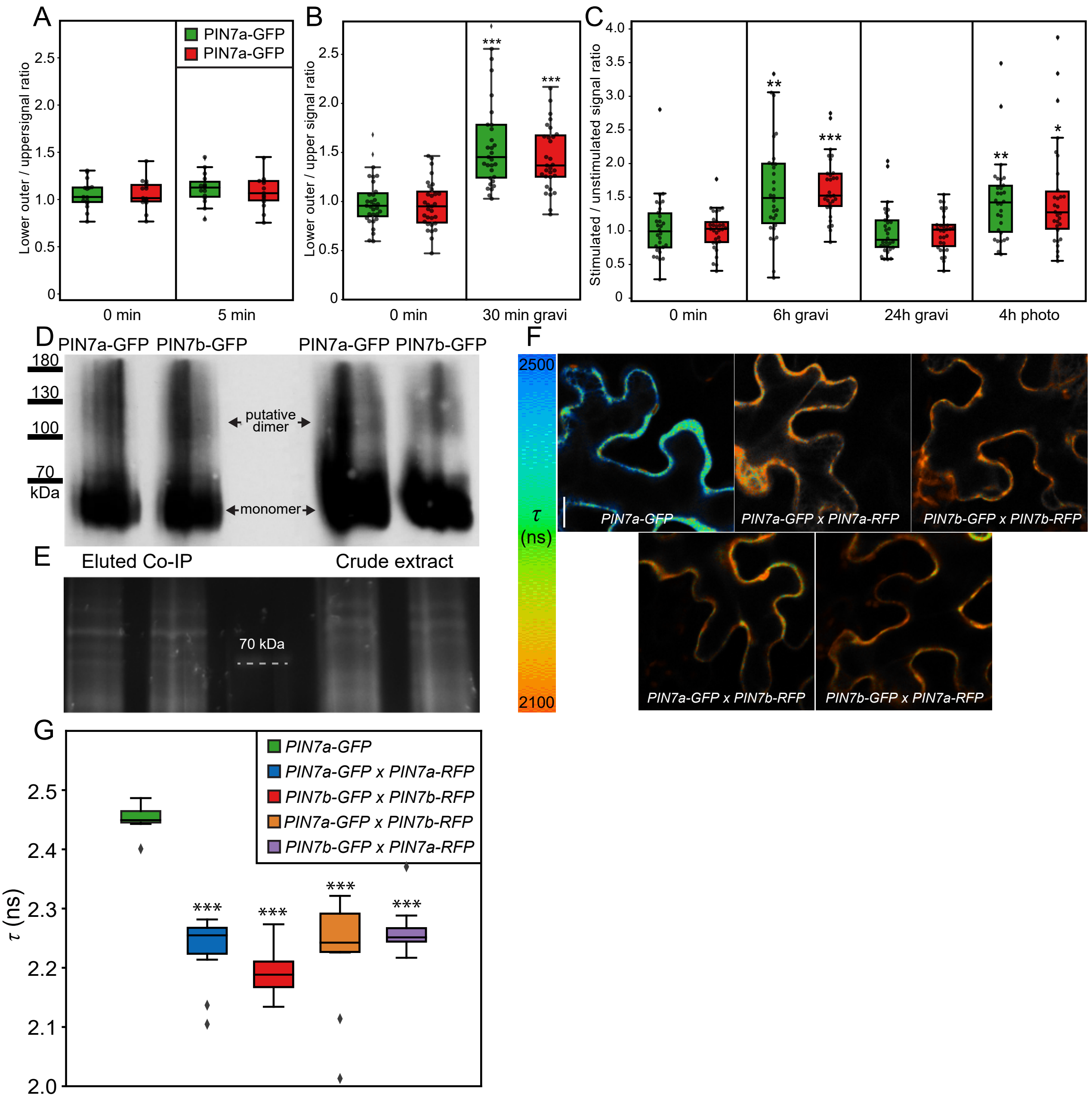


**Supplemental Figure 4.** Polarity change in response to the gravitropic and phototropic stimuli and additional interaction tests of PIN7 isoforms.

**(A)** and **(B)** Quantification of the polarity change of PIN7a-GFP and PIN7b-RFP to the lower side of the root columella cells 5 min **(A)** and 30 min **(B)** after the gravistimulus.

**(C)** Quantification of the polarity change(s) of PIN7a-GFP and PIN7b-RFP in hypocotyls, following the gravitropic and phototropic stimuli.

**(D)** and **(E)** An immunoblot from the native PAGE loaded with the immunoprecipitation eluate and a crude protein extract from the BY-2 cells carrying the *G10-90::XVE>>AtPIN7a-GFP* and *AtPIN7b-GFP* constructs **(D)**; the gel protein staining before the transfer to the membrane **(E)** serves as a loading control. The transgene was induced with 5 μM β-estradiol.

**(F)** and **(G)** The GFP fluorescence lifetime ($\tau$) of tagged PIN7a and PIN7b proteins simultaneously expressed under control of the *G10-90::XVE* promoter in the tobacco leaf mesophyll, shown as a representative image heat map (**[F]**, quantified on **[G]**).

On **(A)**, **(C)** and **(E)**, the middle line represents median, the box 25% and 75% quantiles, whiskers the minima and maxima, dots are single data points, asterisks indicate the difference between the stimulated line and the non-stimulated control (**[A]** to **[C]**), and between $\tau$ of PIN7-GFP in the co-transformed lines and the control on **(G)** (**P* < 0.05, ***P* < 0.01, ****P* < 0.001 by two-way ANOVA). For each line on **(A)** to **(C)**, 15 seedlings and, on **(F)**, **(G)** and **(H)**, 10 cells were analyzed. Bar, 10 μm on **(F)**.
