## Supplemental Figure 5 for "Mutually opposing activity of PIN7 splicing isoforms is required for auxin-mediated tropic responses in *Arabidopsis thaliana*"

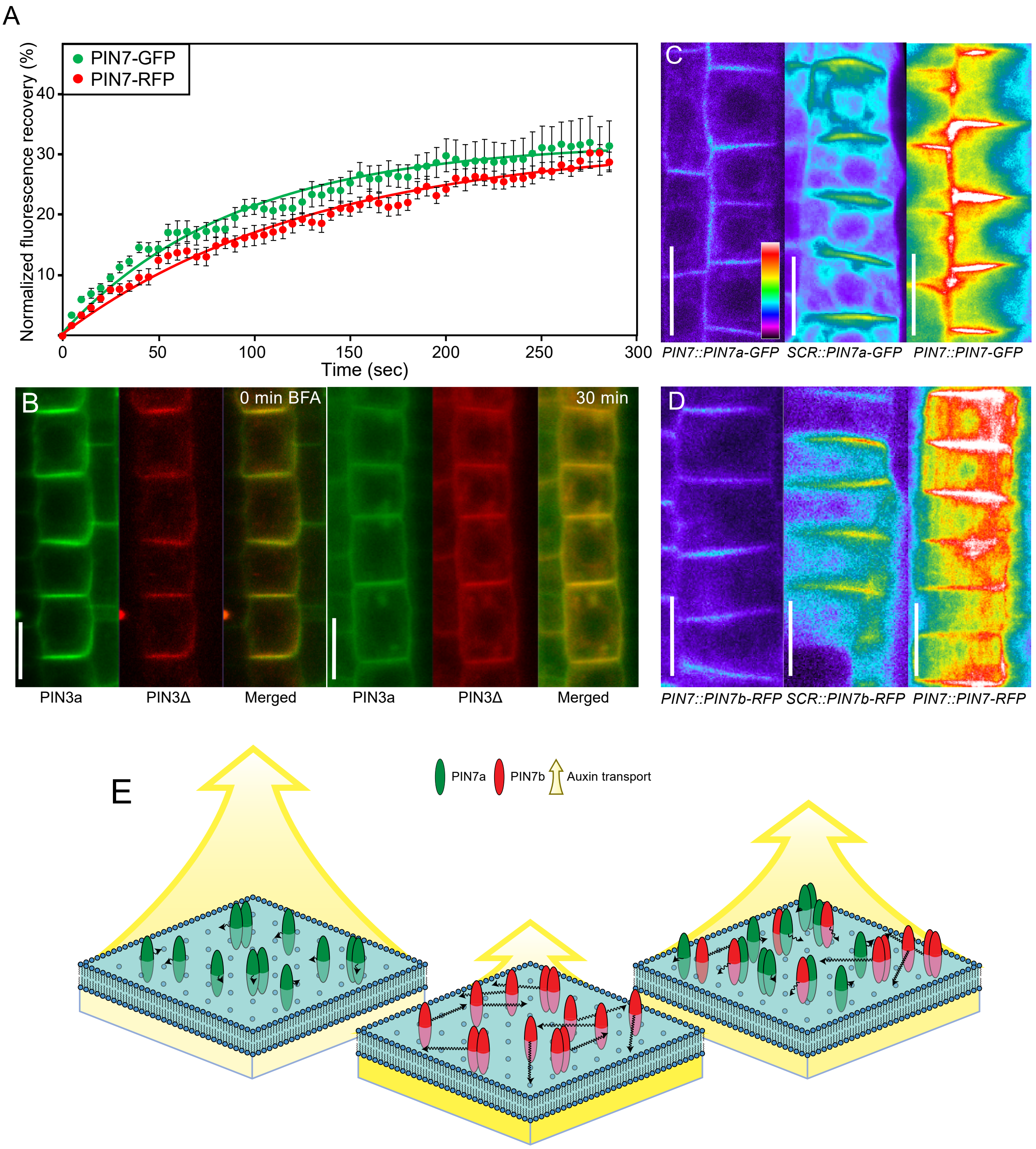


**Supplemental Figure 5.** Additional experiments to the FRAP analysis of the PIN7 isoforms on the plasma membrane.

**(A)** A FRAP on PM in the root tips of seedlings expressing genomic DNA-based *PIN7-GFP* and *PIN7-RFP* transgenes. The data points were fit with the mono-exponential curve. Data are means ± S. E.

**(B)** Comparison of the subcellular localization and response to BFA of the PIN3Δ-RFP protein (designed to mimic the PIN7b isoform), compared to the control PIN3-GFP.

**(C)** and **(D)** A heat map-colored fluorescent intensity of the lines carrying the *PIN7::PIN7a-GFP* cDNA, *SCR::PIN7a-GFP* cDNA and the *PIN7::PIN7-GFP* genomic DNA-based constructs **(C)**, the analogous experimental setup with the lines harboring the RFP is shown on **(D)**. The images were acquired in the root tip pericycle cells with the same microscope settings.

**(E)** Proposed scheme of the interplay between PIN7a and PIN7b and its effect on polar auxin transport in *Arabidopsis*. PIN7a, as monomer or homodimer, occupies the regions with a decreased lateral mobility within PM and mediates polar auxin flow. More diffusible PIN7b interferes with the presence of PIN7a in these regions by its heterodimerization.

On **(A)**, 4 ROIs in each of 5 root tips and, on **(B)** to **(D)**, 3 images in each of 4 root tips were analyzed. Bars, 10 μm on **(B)**, 25 μm on **(C)** and **(D)**.
