## Supplemental Table 1 for "Mutually opposing activity of PIN7 splicing isoforms is required for auxin-mediated tropic responses in *Arabidopsis thaliana*"

| Part | *PIN4a* | *PIN4b* | *PIN4c* | *PIN4a* | *PIN4b* | *PIN4c* | References |
|---|---|---|---|---|---|---|---|
| Aerial | 299 | 314 | 27 | 46,72% | 49,06% | 4,22% | (Cheng et al., 2017) |
| Dark grown seedling | 437 | 333 | 28 | 54,76% | 41,73% | 3,51% | (Cheng et al., 2017) |
| Leaf | 1274 | 1197 | 79 | 49,96% | 46,94% | 3,10% | (Cheng et al., 2017) |
| Light grown seedling | 1463 | 1341 | 52 | 51,23% | 46,95% | 1,82% | (Cheng et al., 2017) |
| Root | 101 | 50 | 4 | 65,16% | 32,26% | 2,58% | (Cheng et al., 2017) |
| Root tip | 167 | 120 | 6 | 57,00% | 40,96% | 2,05% | (Ruzicka et al., 2017) |
| Root apical meristem | 51 | 53 | 19 | 41,46% | 43,09% | 15,45% | (Cheng et al., 2017) |
| Seedling hypocotyl | 41 | 30 | 6 | 53,25% | 38,96% | 7,79% | (Klepikova et al., 2016) |
