## Supplemental Table 2 for "Mutually opposing activity of PIN7 splicing isoforms is required for auxin-mediated tropic responses in *Arabidopsis thaliana*"

| | Name | Sequence |
|---|---|---|
| | BP cloning of natural promoters | |
| 1 | Pr3_GWF | GGGGACAAGTTTGTACAAAAAAGCAGGCTTAtacagcaacactaagtcacaagaa |
| 2 | Pr3_GWR | GGGGACAACTTTTGTATACAAAGTTGTcttgaagggacaaaaatggaaaaccg |
| 3 | Pr4_GWF | GGGGACAAGTTTGTACAAAAAAGCAGGCTTAtctttccttttttgttggttatgtccaaa |
| 4 | Pr4_GWR | GGGGACAACTTTTGTATACAAAGTTGTtttttccggtgggttttggagtt |
| 5 | Pr7_GWF | GGGGACAAGTTTGTACAAAAAAGCAGGCTTActataattttttgtatcggtccaatat |
| 6 | Pr7_GWR | GGGGACAACTTTTGTATACAAAGTTGTattgttgttcgccggagtgg |
| | BP cloning of cDNA constructs | |
| 7 | P3_GWF | GGGGACAACTTTGTATACAAAAGTTGTAATGATCTCATGGCACGACCTCTA |
| 8 | P3_GWR | GGGGACCACTTTGTACAAGAAAGCTGGGTTTTATAACCCGAG |
| 9 | P4_GWF | GGGGACAACTTTGTATACAAAAGTTGTAatgattacgtggcacgacttgtac |
| 10 | P4_GWR | GGGGACCACTTTGTACAAGAAAGCTGGGTTTCAAAGGCCAAGAAGAATAT |
| 11 | P7_GWF | GGGGACAACTTTGTATACAAAAGTTGTAatgatcacatggcacgacctct |
| 12 | P7_GWR | GGGGACCACTTTGTACAAGAAAGCTGGGTTttatagcccgagtaaaatgtagtaaa |
| | BP cloning of SCR promoters | |
| 13 | prSCR_GW_F | GGGGACAAGTTTGTACAAAAAAGCAGGCTTAAATTTTGAATCCATTCTCAA |
| 14 | prSCR_GW_R | GGGGACAACTTTTGTATACAAAGTTGTggagattgaagggttgttggtcg |
| | Primers for inverse PCR of cDNAs | |
| 15 | P4_inv_F | AAATCTAGAGCATATGCCGCCGACAAGT |
| 16 | P4_inv_R | AAACCCGGGTGTTCCGTTGTTGCCGCC |
| 17 | P7_inv_F | AAATCTAGAGCATATGCCACCAGCGAGT |
| 18 | P7_inv_R | AAACCATGGCTTTTACCGGTACAGTTTC |
| | RFP and GFP primers | |
| 19 | RFP_F | AAACCATGGatggcctcctcc |
| 20 | RFP_R | AAATCTAGAtgagtggcggcc |
| 21 | GFP_F | aaaCCATGGGTAAAGGAGAAG |
| 22 | GFP_R | aaaTCTAGAGGATCCATGATG |
| | AS reporter cloning | |
| 23 | P7shift_2F | GATCATCTCAAAATGGTGAAAAACAAAGG |
| 24 | P7shift_2R | CCTTTGTTTTTCACCATTTTGAGATGATC |
| 25 | P7_mutF | GATCTCTGATCATACTCAAAATGGTTAAAACAAAGGTTTTTAAC |
| 26 | P7_mutR | GTTAAAAACCTTTGTTTTAACCATTTTGAGTATGATCAGAGATC |
| | Cloning into pMDC7 inducible system | |
| 27 | PIN4.1codF | ggggacaagtttgtacaaaaaagcaggctctATGATTACGTGGCACGACTTGTACACC |
| 28 | PIN4.1codR | ggggaccactttgtacaagaaagctgggttAAGGCCAAGAAGAATATAGTAGAC |
| 29 | PIN7.1codF | ggggacaagtttgtacaaaaaagcaggctctATGATCACATGGCACGACCTCTACACC |
| 30 | PIN7.1codR | ggggaccactttgtacaagaaagctgggttTAGCCCGAGTAAAATGTAGTAAAC |
